# Functional dynamics underlying near-threshold perception of facial emotions: a magnetoencephalography investigation

**DOI:** 10.1101/383315

**Authors:** Diljit Singh Kajal, Chiara Fioravanti, Adham Elshahabi, Sergio Ruiz, Ranganatha Sitaram, Christoph Braun

## Abstract

Conscious perception of emotional valence of faces has been proposed to involve top-down and bottom-up information processing. Yet, the underlying neuronal mechanisms of these two processes and the implementation of their cooperation is still unclear. We hypothesized that the networks activated during the interaction of top-down and bottom-up processes are the key substrates responsible for perception. We assessed the participation of neural networks involved in conscious perception of emotional stimuli near the perceptual threshold using a visual-backward-masking paradigm in 12 healthy individuals using magnetoencephalography. Providing visual stimulation near the perceptual threshold enabled us to compare correctly and incorrectly recognized facial emotions and assess differences in top-down modulation for these stimuli using coherence analysis. We found a fronto-parietal network oscillating in the lower gamma band and exerting top-down control as determined by the causality measure of phase slope index. We demonstrated that correct recognition of facial emotions involved high-beta and low-gamma activity in parietal networks, Incorrect recognition was associated with enhanced coupling in the gamma band between left frontal and right parietal regions. Our results indicate that fronto-parietal control of the perception of emotional face stimuli relies on the right-hemispheric dominance of synchronized gamma band activity.

## Introduction

Growing evidence hints towards the integration of bottom-up, sensory driven signals and top-down mediated cognitive control playing a vital role in the perception of emotional stimuli (Breitmeyer and others 2006; Ghazanfar and Schroeder; Green and others 2006; Park and Friston 2013). Bottom-up processes that comprise various low-level processes. e.g., visual feature extraction of color, shape, size and orientation (McCarthy and Warrington 2013), characterize the emotion-relevant aspects (emotion-as-stimulus-property) of the presented stimuli (Brozoski and others 1979; Northoff and others 2006). Mapping of neural correlates of the bottom-up processes suggests the active participation of amygdala and hypothalamus in addition to the visual system that encode the affective properties of the presented visual emotional stimuli (McRae and others 2012; Ochsner and others 2009). Visual pathways also contribute to the bottom-up processing via the ventral portion of the striatum, hypothalamic and brain stem nuclei update changes in information such as mismatch, disruption of information flow (Adolphs 2002; Dosenbach and others 2008). Thus, the function of the bottom-up processing comprises reception, registration, awareness of the visual information, and eventually updating if there is any change in the registered information. Top-down processes come into play at the level of processing of the emotional stimulus content (Engel and others 2001; Sarter and others 2001). It is currently understood that although emotional stimuli are registered in the brain by the bottom up processes, the emotional meaning of the stimuli results from top-down processes (Awh and others 2012; Chiesa and others 2013; Mechelli and others 2004; Ochsner and others 2004; Sarter and others 2001). The top-down generation of the emotional response is based on the integration of affective properties and situational-cues (past experience, skill and memory) thus enabling an individual to comprehend the occurrence of a stimulus with emotional properties (Bar and others 2006).

The mechanisms involved in top-down processing of the registered affective properties suggests the formation of recurrent long-distance interactions (Dehaene and others 1998; Dehaene and Naccache 2001; Tononi and others 1998). These interactions involve thalamo-cortical loops, especially activating the prefrontal cortex and higher cortical areas that are inundated by self-amplifying reverberance of network activity. In addition to the above described mechanism, it has also been suggested that attaining a consciously reportable state involves the broadcasting of the affective properties to many functionally specialized brain regions, including those for verbal or motor report (McRae and others 2012; Zanto and others 2011). This broadcasting is performed in a selective way and might adopt re-routing and/or bypassing certain networks and avoiding the visual cortical network (Pessoa and Adolphs 2010). Processing of emotional stimuli in the brain involves rapid modulations of top-down processes to initiate appropriate behavioural outcomes. A plethora of studies suggest that there are two different networks enhancing the top-down mediation of processing an emotional stimulus: fronto-parietal and cingulo-opercular networks. Fronto-parietal networks have been suggested to initiate and adjust the control of the information propagation while the cingulo-opercular networks maintain information until they are over-written or disturbed by another information whichever appears earlier in time (Dosenbach and others 2008).

Top-down processing has been extensively studied using visual backward masking (ViBM) tasks (Dehaene and others 2006; Del Cul and others 2006). In the ViBM paradigm, a prime stimulus is flashed for a brief period (e.f., 16.7 ms) followed by a masking stimulus after a predefined temporal delay. The prime stimulus is not consciously correctly recognized until the temporal delay between the prime and mask exceeds the perceptual threshold of the participant. When the temporal delay between the prime and mask stimuli does not exceed the perceptual threshold information about the prime stimulus can still propagate through the bottom-up processors but will be unable to initiate the subsequent top-down processes, because the propagation of the prime stimulus to conscious processing is disturbed or disrupted by the following mask stimulus and thus will not be supported by the reverberant self-amplifying loops. This defined role of the reverberant self-amplifying fronto-parietal loops are the basis for the global neuronal workspace model (Dehaene and Changeux 2003; Dehaene and others 1998; Dehaene and Naccache 2001; Ramachandran and Cobb 1995).

Any behavioral outcome of top-down and/or bottom up processes results from the coordinated activity in disparate brain regions and specialized networks. This coordinated activity is assumed to be reflected in synchronized activity between distributed brain regions. Neural synchronization is studied by simultaneously measuring the activity from different locations, and assessing if activities at these locations change in a correlated manner. The strength and polarity of the phase synchrony of the oscillatory activity can be interpreted as a proxy for the underlying local neural synchrony, and changes of phase differences between signals analyzed as a function of the frequencies that index the direction of information flow (Ahlswede and others 2000; Nolte and others 2004; Nolte and others 2008a). Several human and primate studies have reported the occurrence of different brain oscillations in supporting the top-down processes (Corbetta and Shulman 2002; Rossi and others 2009; Ungerleider and others 1989; Ungerleider 2000; Webster and others 1994). Del cul and colleagues suggested that for visual perception of stimuli in the prime-delay-target trial, the parietal cortex and frontal regions work in close collaboration (Del Cul and others 2006; Di Lollo and others 2000; Enns and Di 2000; Vorberg and others 2003). Previous work has shown that fronto-parietal networks are involved in the perception of the emotion (Blonder and others 1991; Van Rijn and others 2005), but it is unclear whether the interaction is realized through neuronal oscillatory synchronization or by any other process. Given that top-down processes rely on neuronal synchronization, the direction of information flow becomes relevant: does the synchronization directly reflect top-down information flow, or does it rather reveal the relaying of information to frontal and prefrontal brain regions, or both? We believe that the direction of information flow is crucial for the understanding of the interaction between top-down and bottom-up processes.

In our study, we hypothesize that oscillatory synchronization of long range neural networks is the mechanism by which the emotional content of a stimulus is consciously detected. To rule out any interference from low-level stimulus features we wanted to compare the neural synchronization near the perceptual threshold, i. e., for a temporal delay that results in 70.7% performance level of correct recognition of the emotional face expressions (Leek 2001a; Levitt 1971; Levitt 1992).

To study the long-range network connections involved in the perception of the emotional stimuli, we have chosen a hierarchical approach particularly focusing on the frontoparietal network on a rough spatial scale by defining 4 parcels, i. e., left and right frontal and parietal cortex. In the post-hoc analysis, we have characterized the within and between parcel networks in more detail, by identifying the relevant nodes (brain regions) and edges (strength of functional coupling operationalized as the imaginary part of the coherence) as well as the direction of information flow using phase slope index (PSI) (Nolte and others 2004; Nolte and others 2008a). We then compare the differences in the quantitative strength of the activated brain networks below and above the percetual threshold for processing emotional stimuli using graph theoretic measures.

## Materials and Methods

### 2.1 Participants

Twelve healthy subjects (M±SD=25.5±3.5years, 7 females and 5 males) with normal or corrected-to-normal vision participated in the study. None of the participants had a history of neurological or psychiatric disorders. The study was approved by the local ethical committee of the faculty of medicine, University of Tübingen, Germany. A written informed consent in accordance with the Declaration of Helsinki (Carlson and others 2004) was obtained from all subjects prior to the experiment. All subjects received monetary compensation of 10 Euros/hour for their participation.

### 2.2 Design and Procedure

The experiment consisted of 5 identical runs with each run containing 80 trials and lasting for 8 min, each. A ViBM paradigm was used to manipulate the detectability of the facial emotions by varying the delay between a briefly presented face stimulus and a subsequent mask. We used the adaptive method for choosing the delay for the next trial dependent on the response of the current trial to visually stimulate participants at the threshold for perceiving emotional face expressions. Stimulating near the threshold provides a comparable number of trials with correctly and incorrectly recognized emotional face expressions with identical physical stimulus properties. To study the cortical networks necessary for correctly identifying emotional face expression, brain oscillatory activities were recorded using a whole head MEG.

In our ViBM paradigm, an emotional face either with a positive or negative emotional expression was presented briefly as the prime stimulus (16.7ms), followed by a variable delay (in the range of 0-150ms), and an emotionally neutral mask (250ms) (Fig 1). Evidently, the correct perception of the face expression becomes easier with increasing delays. Each trial started with the presentation of a fixation cross in the middle of the presentation screen. The fixation cross was flashed for 2 sec. Then the prime stimulus (either a happy or sad face) was presented for 16.7 ms (*t_prime_*). The emotional faces were presented in pseudo-random manner across trials. The prime stimulus was followed by a mask stimulus after a variable delay. The delay between the prime and the mask stimuli (*t_delay_*) could be either 0 ms, 16.7 ms, 33.3 ms, 50.0 ms, 66.7 ms, 83.3 ms, 100.0 ms, 116.7 ms, 133.3 ms, or 150.0 ms. A black screen was presented during the delay period. The mask stimulus was flashed on the screen for 250 ms (*t_mask_*). It consisted of a face picture of the same individual as that of the prime, yet with an emotionally neutral expression. Colored face images of both the emotional and the neutral faces were taken from the NimStim Face Stimulus Set (Tottenham and others 2009). 50% male and 50% female faces were selected. The mask was followed by a black screen. The duration of the black screen (*t_blackscreen_*) was chosen such that the stimulation duration of all the trials was of equal length [*t_blacckscreen_* =^1500^ *ms* –(*t_prime_* + *t_delay_* + *t_mask_*)] Thereafter, the instruction cue appeared on the screen prompting subjects to report their valence judgement for the prime stimulus. Participants were requested to report their judgements with their index finger by pressing optical buttons provided for each hand. Response options were presented on the screen as visual cues ‘”NEG+POS’ and “POS+NEG’’, with ‘+’ serving as the fixation cross. The cue “NEG+POS” instructed participants to press the left button if the facial expression of the prime stimulus was recognized as negative and the right button if it was recognized as positive. The cue “POS+NEG” indicated to press the left button for positive and the right one for negative judgements. To minimize any response bias, the two types of cues varied randomly from trial to trial. All participants were requested to respond in each trial and even guess when they were not sure about the valence of the prime face. The response terminated the trial, and the next trial followed thereon. The response interval during which responses were accepted lasted no longer than 2100 ms. The maximum duration of a single trial was 4.5 sec and the inter-trial interval was 5 sec.

**Figure 1:**
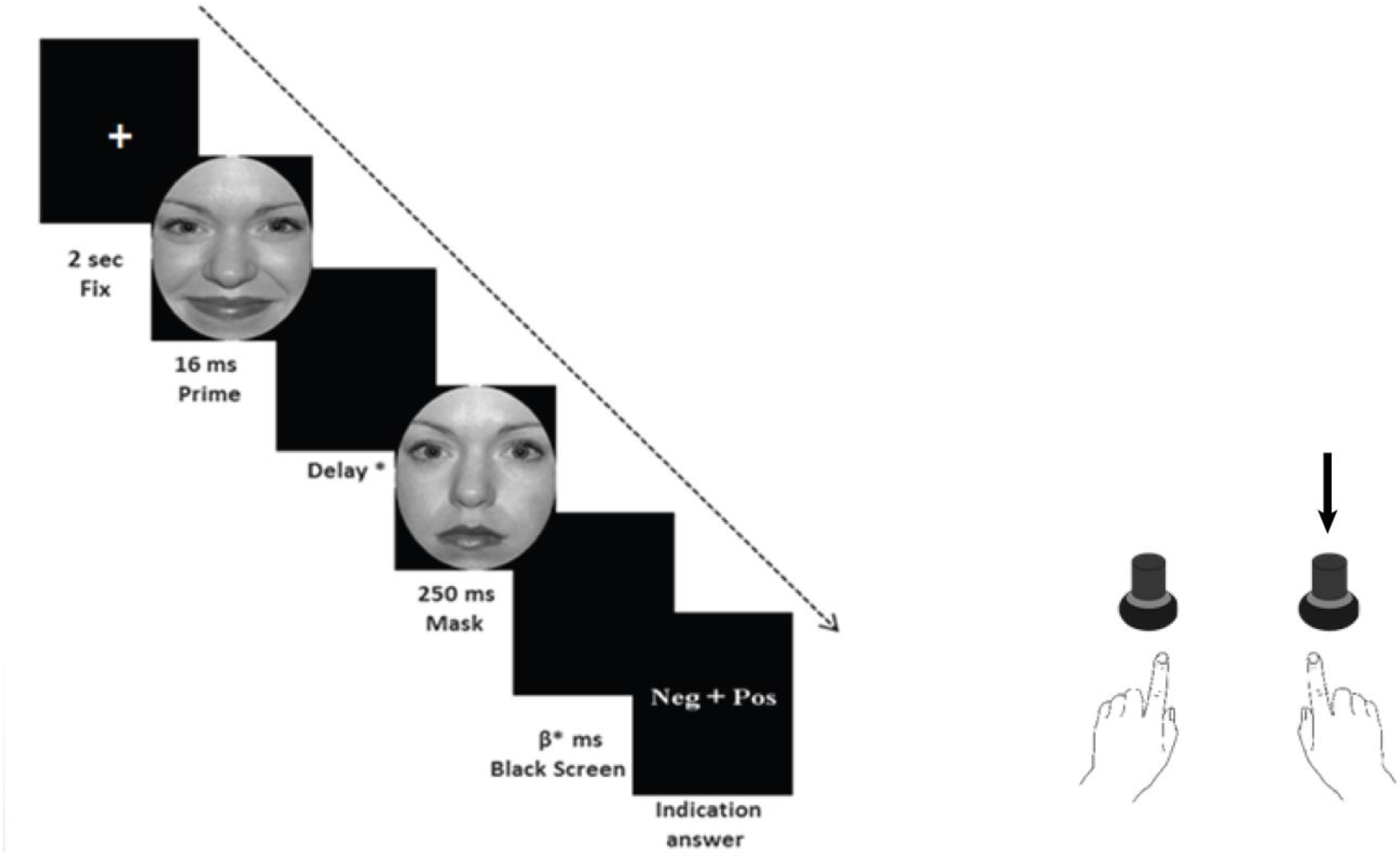
Experimental paradigm: After the presentation of a fixation cross (2 sec), the prime stimulus (an emotionally positive or negative face) was presented for 16.7 ms. After a variable delay (Delay *: 0.0 ms, 16.7 ms, 33.3 ms, 50.0 ms, 66.7 ms, 83.3 ms, 100.0 ms, 116.7 ms, 133.3 ms, and 150.0 ms.), the mask (neutral face) was shown for 250 ms. The black screen, presented for duration Tblank (β *: [1500 ms – (Tprime + Tdelay+ Tmask)]), was followed by the response instructions. Participants were asked to compare the prime stimulus with the mask and to press the right- or left-hand button in order to indicate the emotional valence of the prime. In the above described trial, if the subject correctly identifies the prime face, then the button marked with arrow should be pressed to record the correct response, otherwise, if the other button (not the arrow marked) is pressed, then the response the will be recorded as the incorrectly recognized trial.

To stimulate at the NPT, we used an adaptive procedure (AM) (Leek 2001b; Treutwein 1995). In the AM, the temporal delay between prime and the mask of the next trial is determined based on the stimuli and responses of previous trials. In the AM of our experiment we implemented the ‘two-down-one-up’ rule (TDOU). The rule implies that after any two correct responses the temporal delay between prime and mask becomes shorter by one frame (16.7ms), thus making the task of detecting the emotional expression of the prime face stimulus more difficult. Notably, the two correct responses do not need to be in a row. When the participant responds with an incorrect answer, the temporal delay between the prime and the mask is immediately increased by one frame (16.7ms), thus easing the task in the next trial. Assuming a stationary threshold, the temporal delay is expected to asymptotically reach the threshold for detecting emotional face expressions. This rule converges towards a threshold performance of 66.7% correct.

### 2.3 MEG Recording and Stimuli

MEG (CTF System Inc, Vancouver, Canada) data was acquired using a whole-head 275-axial gradiometer system with a baseline of 5 cm. The MEG is in a shielded room (VaccumSchmelze, Hanau Germany) at the University Clinic of Tübingen, Germany. Brain magnetic data were sampled at the rate of 1072 Hz with an anti-aliasing lowpass filter of 208 Hz. The relative head position with respect to the magnetic field sensors was recorded continuously using three localization coils that were affixed to the left and right preauricular point and the nasion.

Emotional stimuli were presented using an in-house Pascal based program under Dos 6.2 and synchronized with the vertical refresh rate (60Hz) of a screen. The video output of the stimulation computer was sent to a JVC DLA-SX21 projector to flash the stimuli via a mirror system on a screen in the magnetically shielded, dimly lit room. The screen was placed in front of participants in a viewing distance of 70 cm. Stimuli subtended a horizontal visual angle of ~2.5° Participants’ judgements of the valence of stimuli were recorded using in-house built optical buttons. During stimulus presentation, participants were advised to sit still and avoid blinking as much as possible.

### 2.4 MEG data Analysis

Neuromagnetic data were analyzed using in-house MATLAB scripts (MATLAB 2017a) and fieldtrip functions (Oostenveld and others 2011). Data visualization was performed using the BrainNet visualization toolbox (Xia and others 2013). The analysis comprised of the following steps (Fig. 2):

**Figure 2.**
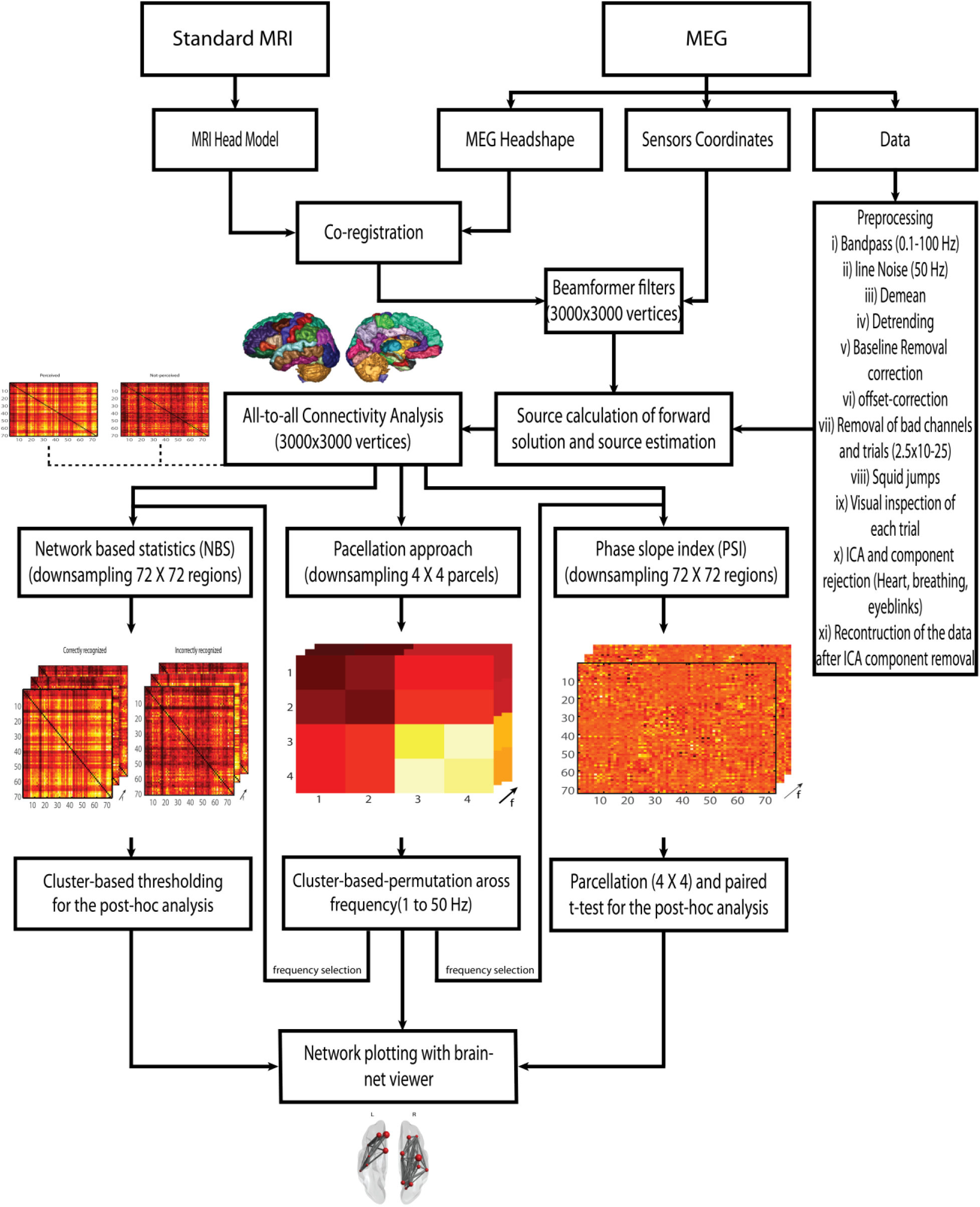
Detailed processing pipeline for the analysis of the MEG data.

#### a Data pre-processing and Artifact removal

Cleaning of the MEG data involved demeaning, detrending, 50 Hz line noise removal and high pass filtering (1 Hz). Magnetic brain data were then inspected visually and trials with large variance across channels and samples (2.5×10^−5^), and abnormal amplitudes were discarded from further analysis. Furthermore, trials containing muscle artifacts indicated by strong broad-band activity, squid jumps, and other non-stereotypical sources were removed. MEG channels with a noise level > 10 fT were excluded from the analysis. Eliminated trials in the MEG artifact rejection were also excluded from the analysis of the motor response. Furthermore, independent component analysis (ICA) was used to remove the contaminating ocular (eye movements, eye blinks) and cardiac artifacts. The ICA using the infomax ICA algorithm (Amari and others 1997; Bell and Sejnowski 1995) decomposed the preprocessed and cleaned data into 100 components. The topography and the waveform of all components were plotted and visually inspected. The components containing eyeblink, eye-movement as well as heart beat and muscular artifacts were removed. From the remaining components, a cleaned MEG signal was reconstructed. Channels containing strong artifacts were marked as bad and excluded from further analyses.

#### b Categorization of the data: Correct and In-correct trials

The preprocessed and cleaned data of the NPT-trials were pooled according to the participants’ valence ratings into: correct trials (trial in which participants correctly identified the prime face) and incorrect trials (trials in which participants were not able to identify the correct emotion of the prime). Due to the two-up-one down rule of the adaptive threshold procedure, there were unequal number of trials for correctly identified (66.7% of trials) and incorrectly identified (33.3% of trials) emotions. We balanced the trial number between conditions by setting it equal to that of the condition with the least number of trials (Nmin), and then by randomly selecting Nmin trials from the condition with the higher number of trials. Frequency analysis was done for the frequency range of 1 to 49 Hz using multi-tapering sliding window fast Fourier transform using discrete prolate spheroidal sequence (DPSS) tapers (Percival and Walden 1993) in steps of 2 Hz. Frequency analysis was done separately for all the trials of correct trials, incorrect trials, and baseline trials.

#### c Condition and frequency specific spatial filter estimation using DICS

Sources of neural oscillatory activity were localized using Dynamic Imaging of Coherent Sources (DICS) (Gross and others 2001), an adaptive spatial filtering method for time-frequency data. In an initial step, neural generators of the magnetic brain activity were localized on a template brain. MEG data were coregistered to an MRI template brain via the three fiducials (nasion, and left and right periauricular point). By discretizing the standard MRI template brain into a regular grid with 1 mm resolution, a template source-model was defined representing 3000 source locations. The standard brain template was segmented, and the single shell spherical head-model representing the electrical properties and the geometry of the brain was computed. Based on the head-model for each grid point the respective leadfield was estimated. Using source- and head-model information, the leadfield reflects the projection of activity from a single source to the sensors. Based on the cross-spectral density and the leadfield matrix, frequency specific spatial filters were estimated that describe the projection of sensor-level activity to the source level. A common spatial filter was computed for both concatenated conditions, i. e. correctly and incorrectly recognized facial expressions.

#### d Calculation of source level connectivity

Source level activity was estimated by applying the frequency specific common spatial filters to the sensor level activity for correctly recognized and incorrectly recognized trials. In the next step, voxel-to-voxel functional connectivity was calculated between all voxels in the brain for each frequency bin, separately for correctly and incorrectly recognized conditions. We parcellated the brain as defined by the Montreal neurological institute (MNI) template into 72 regions. The term *parcellation* here refers to the averaging of all the possible connections arising from voxels in a given seed region to the voxels in a target region, for all seed regions and target regions in the brain. To suppress the confounding problem of spatial spreading of the source activity, we only used the absolute imaginary part of coherence for further analyses (Nolte and others 2004).

#### e Parcellation Analysis

To test our hypotheses of the involvement of fronto-parietal functional coupling in the processing of emotional face expressions, and to reduce the problem of multiple comparison we further merged the brain regions into 4 parcels. Parcellation was done using the spatially normalized T1 brain template provided by the MNI (Collins and others 1998). Parcel I and II represent frontal brain regions, and parcel III and IV the parietal regions. Connectivity strength between parcel l and IV is related to the right intra-hemispheric fronto-parietal connections. Parcels II and III represents the left intra-hemispheric fronto-parietal connections, respectively.

We have segregated the voxel-to-voxel connectivity matrix of N × N (N=72) into 4 × 4 matrix for correct and incorrect trials by averaging the connectivity within and across areas. Within and between parcel connectivity resulted in 10 parcel combinations: (I, I), (I, II), (I, III), (I, IV), (II, II), (II, III), (II, IV), (III, III), (III, IV), and (IV, IV)). To quantify the strength of functional connectivity, we performed a cluster-based permutation test across different frequencies using paired t-test for the absolute part of imaginary coherence to compare the connectivity between correctly and incorrectly recognized emotional face expressions. We looked for both positive and negative clusters, i. e. clusters where the connectivity was larger in the recognized than in the incorrectly recognized emotion and vice versa. We performed a total of 2048 permutation tests (2^12^/2: 12 subjects) by flipping the correct and incorrect conditions across all subjects. To determine the statistical significance of the connectivity differences between correctly and incorrectly recognized trials, the experimental result was compared with the permutation based random distribution. The cumulative probability value for the positive cluster and the negative clusters was 0.025. Network visualization was done using Brain-Net Viewer (Xia and others 2013).

#### f Phase slope index

To identify the direction of the information flow within and between different brain parceled areas during the processing of correct and incorrect trials, we studied the phase slope index on 72×72 connectivity matrix. The phase slope index provides information on whether the signal in one brain region is leading or lagging the signal in another region. The polarity of the index will indicate the direction of information flow. We performed post-hoc analysis and estimated the direction of flow of information for correct and incorrect trials for the frequencies identified in the parcellation analysis and for the individual connections followed by sum and paired t-test (Benjamini and Hochberg 1995).

#### g Network Analysis: Graph Theoretic Network Measures

The main advantage of using graph theoretical measures over classical data analysis for MEG and MRI is that the network architecture of the brain is characterized as a comprehensive metric. In our case we used this measure as post-hoc analysis for the results obtained from the cluster-based permutation across different frequencies for the parceled brain. The graph theoretical analysis was based on the absolute value of the imaginary part of coherence stored in the 72 × 72 connectivity matrix. In the network analysis, each of the 72 brain regions is treated as a node and the strength of coupling from one to another region is considered as an edge. We used the shortest pathlength as a measure to understand the nature of the underlying networks. We then assessed the difference in the synchronized network organization for correctly recognized and incorrectly recognized trials across different frequencies. (Benjamini and Hochberg 1995; Benjamini and Yekutieli 2001; 2005).

A 72×72 connectivity matrix for each subject is processed for the correctly recognized and incorrectly recognized networks. The connectivity matrix for correctly recognized and incorrectly recognized trials were randomly mixed, vectorized and sorted in descending manner. Only 20% of the shortest pathlengths of the vectorized network areas were taken for further analysis. A t-test contrasting the two groups (correctly recognized and incorrectly recognized) was then computed for each pairwise association. Any association with a t-statistic exceeding 1 was admitted to the set of suprathreshold links used by the Network Based Statistics toolbox (NBS) (Zalesky and others 2010). The NBS was implemented with 10000 permutations to generate the null distribution of maximal component size. The NBS performs the clustering in the topological space rather than the physical space. The NBS is the graph analogue of the cluster based statistical method.

## Results

### 3.1 Parcellation Approach

The goal of our study was to test the hypothesis of the participation of synchronized fronto-parietal neural network in the processing of emotional face stimuli. To this end, we investigated the differential involvement of the functional connections for correctly or incorrectly recognition of emotions in face stimuli. To avoid the problem of multiple comparisons, we performed parcellation of the brain followed by the cluster-based-permutation across different frequencies (Benjamini and Yekutieli 2001; 2005) for the correction of type I errors.

Cluster based permutation across frequencies revealed a significant difference between correct and incorrect recognition of emotions in face stimuli at 35 Hz (Fig. 3 A) ii), for the absolute imaginary part of the coherence (parcel I and IV: p (11)=0.01, t=4.41). The difference between correctly and incorrectly recognized trials was not significant in other frequency bands. To assess the direction of the information flow in the right hemisphere, we computed the phase slope index for the connections crossing the frontal and parietal parcels and found a significant difference between correctly recognized and incorrectly recognized trials. A significant positive difference between correctly recognized and incorrectly recognized trials of PSI values was found for the right frontal and parietal parcels (parcel I and IV: p(11)=0.01, t=2.87) in the gamma band (35 Hz). Positive PSI values suggest the flow of information from frontal regions to parietal regions. To clarify whether the directed interaction originates from recognized or incorrectly recognized trials, PSI was compared against zero. While the PSI for incorrectly recognized trials (p(11)=0.54, t= 0.63) did not differ from zero, it was significantly different (p(11)=0.04, t= 2.25) for recognized trials at 35 Hz (Fig. 3 A). Further analysis using the parcellation approach demonstrated that the right parietal parcel revealed significantly more within-parcel connections in the recognized than in the incorrectly recognized condition at higher beta (29 Hz) and lower gamma frequencies (31 and 33 Hz). The right parietal cortex (parcel IV) was significantly active as found by the cumulative probability threshold (p(11)=0.02, t=3.6) (Fig. 3 B). Furthermore, we investigated the contribution of interhemispheric connections for the perception of emotional stimuli. We found stronger connectivity for the incorrectly recognized than for the recognized trials between left frontal and right posterior cortex. The cumulative probability distribution in the gamma band (37 to 41 Hz) was found to be (t(11)=-3.54, p=0.02). (Fig. 3 C).

**Figure 3.**
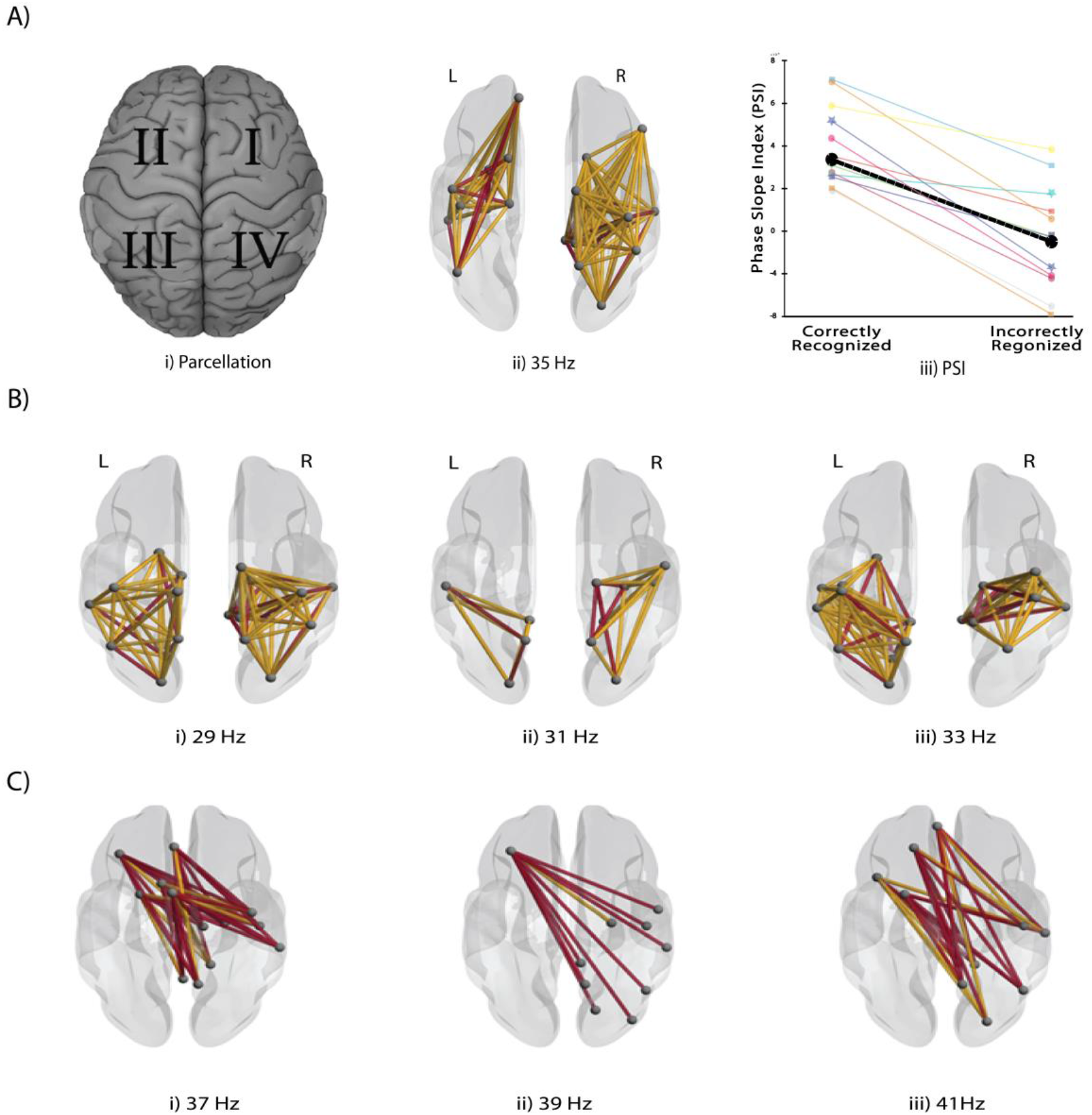
A: i) Partition of the whole brain into four parcels. ii) Parcellation of the right and left hemisphere representing the difference between correctly recognized and incorrectly recognized emotions at 35 Hz. iii) PSI values for correctly recognized and incorrectly recognized trials for the right fronto-parietal network at 35 Hz. B) Difference in network architecture between correctly recognized and incorrectly recognized emotions: i) High beta (29 Hz), ii) and iii) Low gamma (31 and 33) in the right parietal parcel. C) Significant functional interactions between the left frontal parcel and right parietal parcel while contrasting the percieved with incorrectly recognized emotions. i) at 37 Hz, ii) at 39 Hz and iii) at 41 Hz in the low gamma band activity. The positive connections of the network architecture are represented in yellow and negative connections in red.

### 3.2 Post-hoc analysis using Graph Theoretical Measures

Brain networks can be mathematically represented as graphs consisting of a set of nodes (brain regions) and edges (functional or structural connections between brain regions). The pairwise couplings are summarized in the form of a network connection matrix. The graph theoretical measure Cluster Coefficient reflects the local integration, and Shortest Pathlength suggests the level of global integration within the network (Bullmore and Sporns 2009; Rubinov and Sporns 2010; Sporns 2010). We estimated the shortest path using the Floyd-marshal algorithm (Floyd 1962). We studied the network organization using the network-based statistics (NBS) toolbox (Zalesky and others 2010). The post-hoc analysis of the network organization at 35 Hz and estimation of shortest pathlength was conducted using the NBS toolbox for 10000 iterations (Fig. 4 A iii. Positive t-value indicates that the shortest pathlength is shorter for correct trials compare to incorrect trials, and negative t-value indicates that the shortest pathlength is shorter in the incorrect trials (p (11)=0.001, t=3.2) than in the correct trials. We also found significant difference in the correctly and incorrectly recognized trials in the right parietal hemisphere at the following frequencies: 29, 31 and 33 Hz.

**Figure 4.**
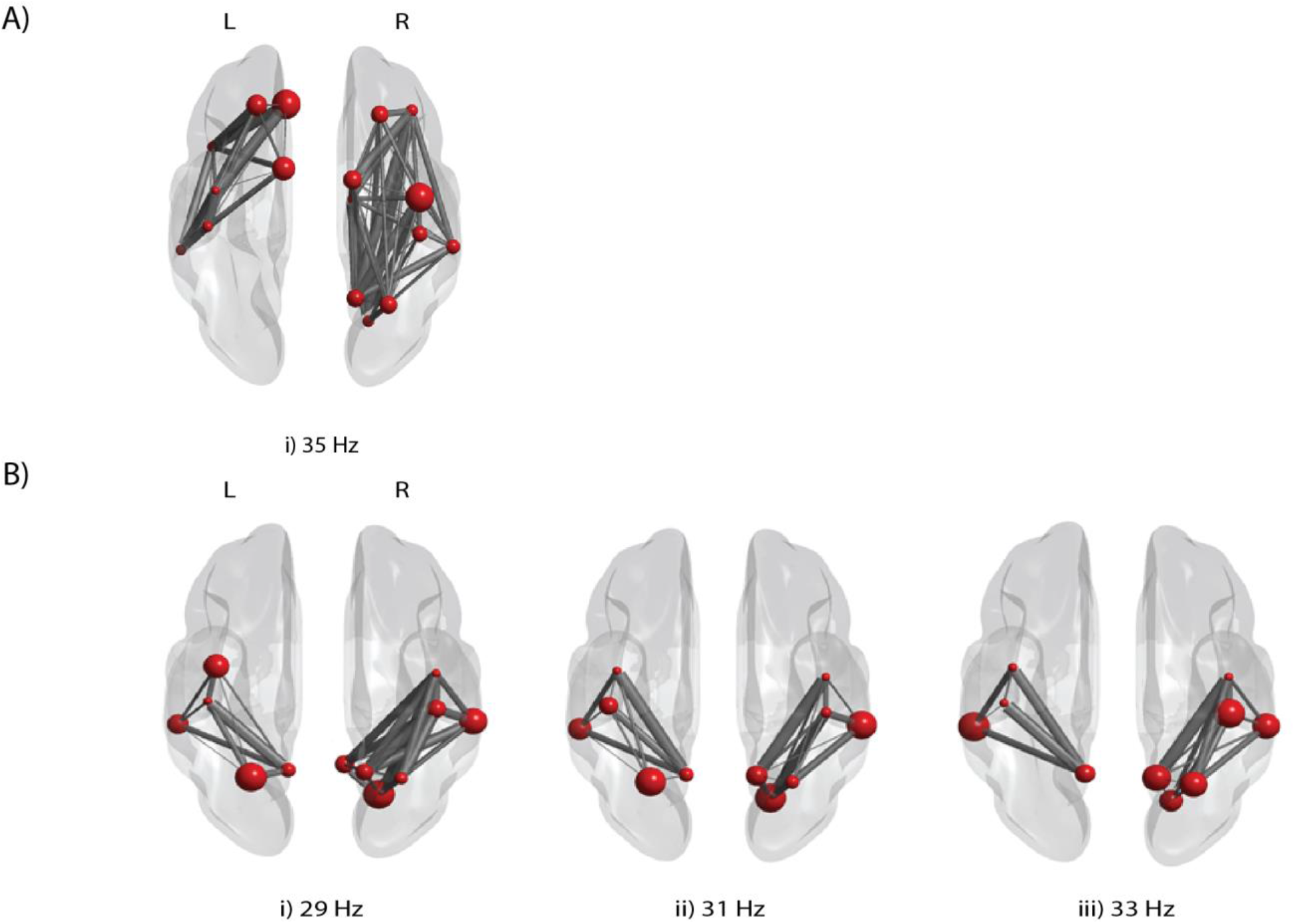
A i) Network organization at 35 Hz. The right hemisphere shows significant difference for the shortest pathlength for correctly recognized and incorrectly recognized trials. B) i), ii) and iii network organization at 29 Hz, 31 Hz and 33 Hz. Thicker lines represent low value for the shortest pathlength.

**Figure 5.**
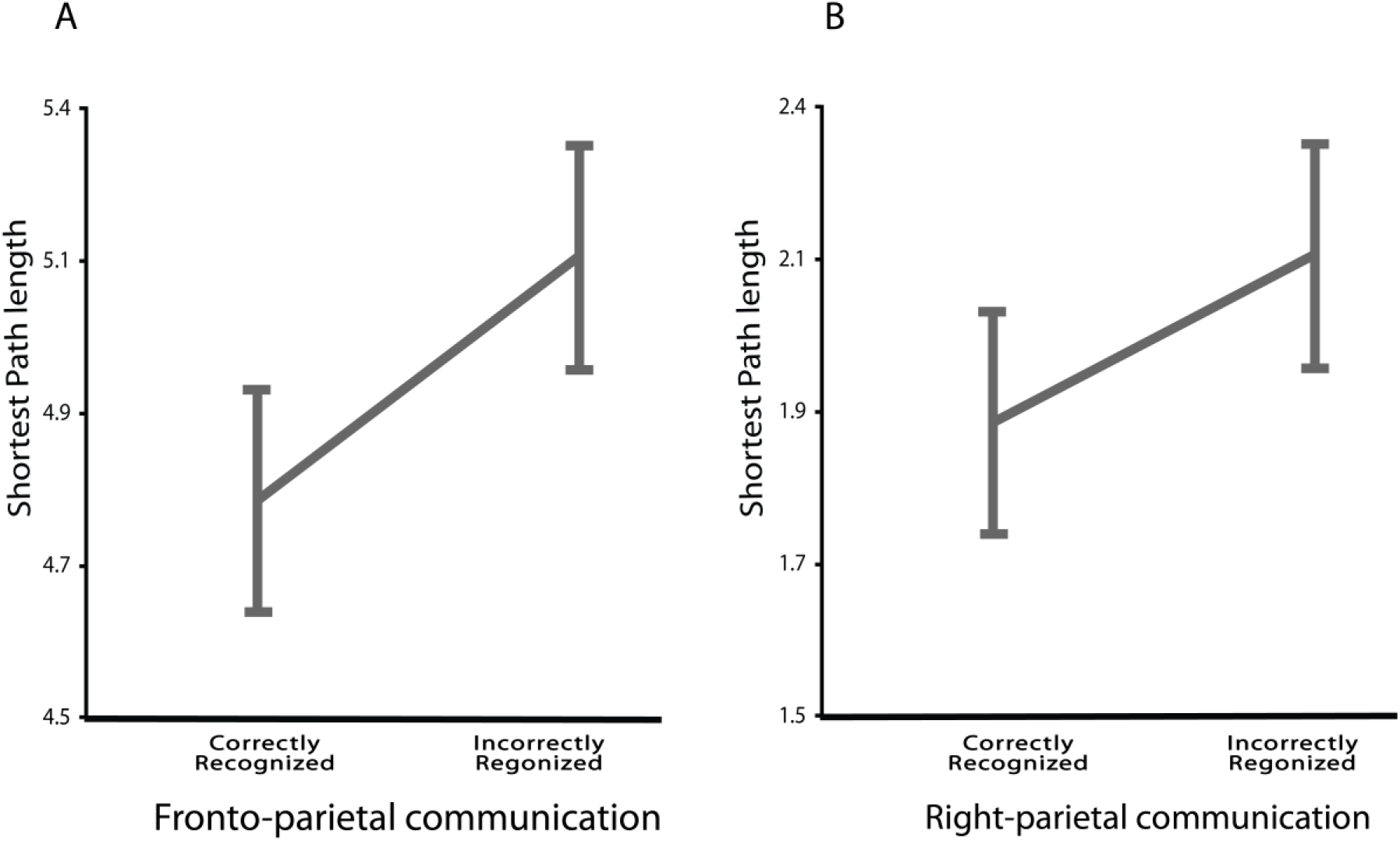
Plots showing the shortest pathlengths for correctly recognized and incorrectly recognized trials. a) Shortest pathlengths at the fronto-parietal interactions at 35 Hz. Correctly recognized (M±SE=4.85±0.064) and incorrectly recognized (M±SE=5.01±0.063) emotions, b) Right parietal interaction for recognized (M±SE=1.88±0.05) and incorrectly recognized ((M±SE=2.12±0.06) emotions.The integer value on the y-axis represents the number of minimum nodes required for an information to reach from point 1 to point 2. In our experiment, it suggest the number of nodes in the fronto-parietal communication and right-parietal communication for correctly recognized and incorrectly recognized trials

## Discussion

ViBM in combination with neurophysiological recordings is an established paradigm to study the neural mechanisms underlying the perception of emotional stimuli. While previous work has highlighted the activation of brain regions that are mandatory for the correct interpretation of the valence of facial expressions (Adolphs 2002; Dolan 2002; Liu and others 2002), here we studied the functional connections between regions, i. e. the neural networks that are involved in the perception of emotional face stimuli. Towards this end, we studied the organization of brain networks using the measure of functional connectivity near the perceptual threshold of human participants. To rule out any physical differences of stimulus parameters between correctly recognized and incorrectly recognized emotional faces accounting for brain activation and network differences, stimuli were presented near the perceptual threshold resulting in correctly recognized and incorrectly recognized trials for the same masking stimulus. Neural networks involved in the processing of emotional face stimuli were identified by contrasting correctly recognized facial emotions to incorrectly recognized emotions. Except for correctly guessed emotional stimuli, correctly identified responses were assumed to reach a conscious level. In contrast, incorrectly recognized and correctly guessed emotions were supposed to reflect rudimentary emotional processing not reaching full awareness. Although in correctly recognized trials correctly guessed and correctly identified emotions cannot be distinguished, there are in any case more consciously correctly recognized emotions than in the incorrectly recognized condition.

Based on the model that the correctly recognized emotional valence relies on the well-adjusted interplay between fast bottom-up and slower top-down processes (Delorme and others 2004), one can conclude that stimulating with a prime-delay-mask near the threshold of emotion perception provides enough time to complete the fast bottom-up processing of feature extraction, while possibly interfering with later top-down processes. Thus, by studying perception of emotions near the perceptual threshold, we would be able to assess the network dynamics of correct and incorrect recognition of emotional stimuli. This interpretation is supported by our findings that do not show activation in the primary visual cortical areas when comparing between correct and incorrect trials.

We have particularly assessed the brain networks involved in top-down processing that is assumed to be maintained by reverberating long-range fronto-parietal network connections. Our results show that there are stronger fronto-parietal functional connections stimuli predominantly on the right hemisphere for correctly recognized emotional face expressions. In contrast, left hemispheric frontal influence on right parietal cortex appears to be detrimental for the analysis of the emotional valence of facial stimuli.

### Parcellation Approach

From the literature (Cahill and others 2004; Canli and others 1998; Silberman and Weingartner 1986; Wager and others 2003), we have the understanding that sensory areas closely interact with frontal as well as parietal areas when they are subjected to the processing of the emotional stimuli. The fronto-parietal interaction most likely involves an extended network spanning multiple regions in the prefrontal, frontal and parietal cortex rather than in the form of a point-to-point connection.

To test the participation of fronto-parietal network in the top-down processing of emotional stimuli, we have adopted an approach using cluster-based permutation of source level connectivity across different frequencies. We demonstrated, firstly, that the fronto-parietal network were significantly more functionally coupled in the gamma band during the recognition of emotional faces in contrast to incorrectly recognized emotional faces. Secondly, using the phase-slope index, we could demonstrate that in these networks the direction of information flow is from right frontal regions towards the right parietal cortex (Adolphs 2002; 2003). Thirdly, within-connections of the right posterior cortex were significantly stronger in high beta and low gamma frequency bands during correctly recognized as compared to incorrectly recognized emotional faces. Fourthly, we showed that there is higher connectivity for incorrectly recognized stimuli than for correctly recognized stimuli between left frontal and right parietal regions in the gamma frequency range.

### Gamma Frequency Band

Fronto-parietal network differed between correctly recognized and incorrectly recognized emotional stimuli mainly in the gamma band. Gamma frequency activity in emotion perception has generally been related to bottom-up sensory processing (Li and Lu 2009; Müller and others 1999). Synchronized neuronal firing in the high frequency range in the human cortex has been suggested to reflect the formation of Hebbian cell assemblies (Eckhorn and others 1990; Pulvermüller and others 1995; Singer and Gray 1995), and can be recorded with E/MEG in sensory processing. Gamma modulation has been reported to occur for the variation of features of the visual stimulus (Müller and others 1996; Müller and others 1997; Tallon-Baudry and others 1997; Tallon and others 1995) and during perception (Keil and others 1999a; Keil and others 1999b; Tallon-Baudry and others 1996; 1997).

Previous studies have investigated oscillatory brain activity in emotion perception with stimuli well above the perceptual threshold. Thus the previously described gamma-band activity might be due to specific emotion processing, and also to low-level visual processing. In contrast to the previous studies, in our study we have stimulated the emotion networks near the perceptual threshold. By comparing correctly recognized and incorrectly recognized emotions we could rule out networks involved in low-level feature extraction and highlight the fronto-parietal top-down control networks oscillating in the gamma frequency range and thus interacting with bottom-up passage of information. The participation of the gamma frequency band is found to be significant in the right parietal cortex which is in line with findings attributing an important role of the right hemisphere in emotion processing(Balconi and Lucchiari 2008; Keil and others 2001; Müller and others 1999). A negative contribution (stronger connectivity for incorrectly recognized stimuli) from the left frontal cortex to parietal cortices was also observed. It seems very plausible that the functional coupling in the gamma band is a proxy for the correct perception of emotional faces in our experiment (Eckhorn, et al., 1990; Pulvermüller, et al., 1995; Singer and Gray, 1995). We argue that the involvement of different networks from within and between hemispheres via gamma oscillations is the basis for the generation and propagation of reverberant self-amplifying fronto-parietal loops (Baumgarten and others 2017; Dehaene and Changeux 2003; Dehaene and others 2014; Dehaene and others 1998; Dehaene and Naccache 2001; Del Cul and others 2006) enabling the perception of emotions.

### Phase Slope Index (PSI)

Using PSI, the direction of information flow from one region to another can be inferred (Nolte and others 2008b). We have estimated the PSI values between individual areas in different parcel combinations across different frequencies identified in the cluster-based permutation parcellation analysis for correctly recognized and incorrectly recognized trials. We found that the PSI value indicates a significantly positive contribution from right frontal cortex (parcel I) toward right parietal cortex (Parcel IV). The positive PSI value between ‘A’ and ‘B’ suggest the activities are leading at ‘A’ and ‘A’ is directing information towards ‘B’. Using this analysis, we were able to demonstrate that the frontal cortex is leading and controlling the information flow towards the parietal cortex. This finding supports the interpretation of fronto-parietal network exerting frontal top-down control over parietal areas.

Except for the right fronto-parietal interaction, we were unable to demonstrate any other significant directed information flow between left frontal cortex and right parietal cortex or any other combination. From this result one might conclude that the communication between left frontal and right parietal cortex is bi-directional and/or may be dominated by noise. However, given the predominant involvement of the right hemisphere in top-down emotional control it might also be argued that information exchange between the other regions is less dominant.

Our finding of the right-hemisphere directed fronto-parietal information flow in the processing of emotions are consistent with previous findings on emotion perception that suggested strong participation of the right hemisphere in the perception of emotion irrespective of the valence of the emotional stimuli (Davidson 1984; Ehrlichman 1987; Hirschman and Safer 1982).

### Positive and Negative emotional faces

It has been documented in the literature that left hemisphere is involved in the processing of positive emotion processing and right hemisphere is responsible for the processing of the negative emotional valence (Aftanas and others 1998; Müller and others 1999; Tucker 1981; Tucker and Dawson 1984). In our experiment, using a ViBM at the NPT, we did not find any significant difference in network oscillations for positive and negative faces.

### Graph Theoretical Network Approach

For the past two decades, various graph theoretical measures had been extensively used to study structural and functional neural architecture of the brain. In our analysis shortest pathlength was differentially expressed in brain networks prevalent for correctly recognized and incorrectly recognized emotional face expressions with shorter average pathlengths for the former condition. In graph-theoretical network analysis nodes correspond to brain areas and edges correspond to the connection between nodes. The shortest pathlength between two nodes is the minimal number of edges to reach one node from one another. The average shortest pathlength for one node is the average of all shortest pathlengths from this node to all other nodes. Mapping the average shortest pathlength for individual brain regions provides insights on how well individual areas are connected to the rest of the brain. A low value of the local average pathlength is an index for the integration of an area in the network.

The pathlength of the network is an important predictor of the network performance (Vragović and others 2006). The performance of a network in basically dependent on the network average pathlength: *‘’The shorter the pathlength, the better the performance”.* However, in the diseased brains such as schizophrenia and epilepsy, due to insufficient inhibition all-to-all connectivity between brain areas is generally increased. Thus although the shortest pathlength is decreased, functional performance is most likely not improved (Andreou and others 2015; Yan and others 2017). In contrast, looking at only the strongest and functionally relevant connections by selecting the shortest pathlengths as we did in our analysis might yield more meaningful results. In our study we demonstrated that the shortest pathlength for the right fronto-parietal subnetwork during the correct perception of emotional face expressions supports the notion of right fronto-parietal communication mediated by oscillatory gamma band activity being crucially important for the processing of emotions.

The human brain consists of the disparate functionally specialized regions, and the information exchange between them is either task dependent and/or default mode. Here, the task dependent communication requires to be very precise and efficient in the integration of the information from different brain regions and can be characterized by the graph-theoretical measures. A key feature of the healthy brain is the optimum balance between segregation and integration of information exchange between brain regions (Tononi and others 1998). For example, the shortest pathlength, a global characteristic, is an index for the functional integration of the brain (Achard and Bullmore 2007), and thus suggests how easy it is to transport information or other entities within the network. Shortest pathlength has been demonstrated to promote the effective integration across cortical regions, implying that, in contrast, relatively longer pathlengths might indicate that the communication between connected regions is slower, with reduced strength of connectivity, and less efficient (Achard and Bullmore 2007; Bassett and Bullmore 2006)

Our results on shortest pathlengths are in line with clinical studies which have shown that tight integration of two region can be studied using the shortest pathlength metric in schizophrenia (Liu and others 2008; Wang and others 2010), autism (Barttfeld and others 2011), and stroke (Adams and others 1993; Crofts and others 2011; Tenenbaum and others 2000). In schizophrenia, task related pathlength was found to be increased in the alpha, beta and gamma frequency bands (Breakspear and others 2006; Micheloyannis and others 2006; Rubinov and others 2009). Our study is in line with Dahaene et al (Dehaene and others 1998; Dehaene and Naccache 2001) proposing the general involvement of the fronto-parietal regions in the perception of masked stimuli. Adding to these findings, our results suggest a highly specific right-hemispheric fronto-parietal network for the successful processing of emotional stimuli.

## Conclusion and future direction

Studies on the processing of emotional stimuli have shown that in schizophrenia, there is a significant increase in the global pathlength compared to the healthy participants (Berkovitch and others 2018; Liu and others 2008). The perceptual threshold of the schizophrenic patient is elevated and thus affects the top-down control of emotional processing. It has been argued that the increase of the threshold for emotional processing makes these patients unable to perceive emotions correctly.

We suggest that the proposed research is highly important for answering fundamental questions about networks involved in conscious and non-conscious perception of the emotional faces, and thus provide the basis for novel treatments of neurologic and psychiatric disorders through electro/magnetic stimulation (Bejjani and others 1999; Frank and others 2007; Romei and others 2012), brain-computer interfaces, and neurofeedback (Birbaumer 2006; Birbaumer and others 2008; Birbaumer and others 2013; Kajal and others 2017; Kajal and others 2015; Sitaram and others 2017; Sitaram and others 2008). Specifically, we suggest exploring new avenues towards actively understanding the neural mechanisms underlying basic perceptual processes in syndromes as different as neglect, schizophrenia and autism.

## Summary

In our study, we found that in the gamma frequency band, the right fronto-parietal network, the parietal network, and the left frontal to right parietal functional connections are significantly coupled during the perception of the emotional stimuli at near perceptual threshold. This gives us the understanding that the perception of emotional face expression involves functional coupling between different areas and is mediated by phase-locked oscillatory activity in the gamma frequency band. Our results support the framework of the global neuronal workspace theory which suggests that directed frontoparietal connections set the coordinated interplay between bottom-up with top-down processing.

## Acknowledgements

We would like to thank Mr Jürgen Dax, for the help in the setting up of stimulation presentation setup. This work was supported by Comisión Nacional de Investigación Científica y Tecnológica de Chile (Conicyt) through Fondo Nacional de Desarrollo Científico y Tecnológico, Fondecyt (project n° 1171320 and 1171313), CONICYT-PIA Anillo ACT172121.

